# Neural and Behavioral Changes in Older Adults from Auditory-Cognitive Training

**DOI:** 10.1101/2025.04.01.646593

**Authors:** Charlie Fisher, I.M. Dushyanthi Karunathilake, Michael A. Johns, Allison Vance, Stefanie E. Kuchinsky, Samira Anderson, Jonathan Z. Simon

**Affiliations:** University of Maryland, College Park, MD 20742 USA; Walter Reed National Military Medical Center, Bethesda, MD 20910 USA

## Abstract

Speech perception in noisy environments is a common challenge among older adults, even for those with clinically normal hearing. Cognitive decline may be one of the contributing factors, and, as such, auditory-cognitive training may enhance speech perception in these conditions. This study aims to determine if auditory-cognitive training can improve speech-in-noise listening in normal-hearing, older adults using neural and behavioral measures, supplemented with comparisons across younger and older adults. Neural responses were obtained using magnetoencephalography (MEG) while participants listened to long, narrative passages (60 s) under four noise conditions. Neural measures employed reverse correlation using encoding and decoding models, via the temporal response function (TRF) framework, to predict neural responses and reconstruct stimulus features, respectively, with the boosting algorithm to enforce sparsity. Behavioral measures, such as working memory (reading span; RSPAN), speech perception in noise (SPIN), and nonlinguistic auditory stream segregation (stochastic figure-ground; SFG) showed improvement post-training, along with neural and subjective ratings for listening effort. Additionally, auditory-cognitive training may enhance the neural contrast between the selectively attended and unattended stimulus reconstructions, and pre-training SFG performance may predict the extent of this neuroplasticity change. These results provide promising, additional insight into the effects of auditory-cognitive training, both perceptually and neurally.

## 1. INTRODUCTION

Auditory stream segregation is the process of listening to a single sound source, e.g. a speaker, among a background of noise or other distractors. As the signal-to-noise ratio (SNR) decreases, this can introduce listening difficulties, regardless of age. Older adults, however, even those with clinically normal hearing thresholds, show greater listening difficulties [1] and report higher listening effort in noisy environments than younger adults [2]. This may be due to age-related declines in auditory and/or cognitive processing [3]; while interventions remain limited, evidence supports training with jointly auditory-cognitive tasks as a possible option [4]. Auditory-cognitive tasks require the engagement of both auditory and cognitive processes, such as in auditory stream segregation. Note that auditory-cognitive tasks stand in contrast to the more common auditory-based cognitive tasks, in which cognitive tasks (e.g., recall) are presented using auditory stimuli, and for which the focus is more cognitive than auditory. Auditory-cognitive training may provide the most benefit for speech-in-noise perception—as it would target the same underlying mechanisms [5], but the neural impact of such training remains largely unexplored.

Beneficial neuroplasticity changes have been shown after auditory-based cognitive training in predominately hearing-impaired, older adults, subcortically [6], [7] and cortically [8]. Structural MRI changes have also been found in predominately hearing-impaired, older adults after an adaptive auditory-based cognitive training [9]. However, no subcortical neuroplasticity changes were observed in relatively normal-hearing older adults with an auditory-based cognitive training [7]. Encouragingly, when training incorporated auditory-cognitive tasks, subcortical neuroplasticity changes were found even with younger, normal-hearing adults [10]. It remains unclear whether these changes would be observed in normal-hearing, older adults undergoing auditory-cognitive training, particularly in the cortex. Here, we address both questions using auditory-cognitive training and analyzing cortical responses in older adults with normal-to-near-normal hearing thresholds. Investigating the neural effects of auditory-cognitive training may provide valuable insight into agerelated speech-in-noise difficulties and the neural mechanisms that training benefits.

Previous research has observed age-related neural differences in the auditory cortex using reverse correlation with encoding and decoding models [11], [12]. These analysis frameworks provide a detailed measure of neural responses during auditory stream segregation tasks. Encoding models can be used to identify the timing and fidelity of time-locked neural responses to speech features, and decoding models can be used to correlate reconstruction stimuli with actual stimuli to measure cortical representation, or information processing. Both methods provide a quantifiable framework to track neuroplasticity changes across time, necessary to see any neural effects of auditory-cognitive training. In this study, we implemented two adaptive auditory-cognitive trainings in older adults with normal-to-near-normal hearing thresholds and used this framework to analyze differences before and after training, and between age groups.

The goal of this magnetoencephalography (MEG) study was twofold: first, confirm the previously seen neural differences between younger and older adults, and then investigate any neural differences in older adults between pre- and post-training. This work shows that auditory-cognitive training, in addition to showing behavioral improvements, also shows positive changes in neural measures. Importantly, we found that attentional contrasts (attended vs. unattended) feature representations were neurally present for younger adults but were not for older adults. After training, however, older adults showed increased attentional contrasts. This suggests that, after training, the neural representation of the attended speaker becomes stronger than that of the unattended speaker, which may reflect a reduction in listening difficulty. Additionally, the level of attentional contrast change is predicted by a behavioral measure of nonlinguistic auditory stream segregation: performance on a stochastic figure-ground (SFG) task. These results provide additional support for auditory-cognitive training in older adults with relatively normal hearing thresholds and highlight that the benefits of training may be predicted by a behavioral task.

## II. Methods

### A. Participants

24 younger adults (18-28 years old) and 23 older adults (60-83 years old) participated in this study. All participants were monolingual, native English speakers. Inclusion criteria for this study included no indication of cognitive impairment and relatively normal hearing, defined as a Montreal Cognitive Assessment (MoCA) score of or above 26 and pure tone hearing thresholds of less than or equal to 25 dB (250-4000 Hz), respectively.

To measure the effects of training, older adults completed pre- and post-training neural and behavioral tasks. Younger adults did not undergo auditory-cognitive training and, therefore, completed a single session of neural and behavioral tasks.

Two older adults were excluded from neural analysis due to head size exceeding MEG limits. One older adult was excluded from session comparison analysis for neural and subjective rating analysis due to a metal artifact in their post-training MEG scan. An additional older adult was removed from listening effort analysis due to not understanding the rating scale. One younger adult was excluded from the SFG task due to data loss.

### B. Auditory-Cognitive Training

Older adults were randomly assigned to one of two training groups with different levels of cognitive demand, but all are combined here for simplicity. Younger adults did not undergo any training.

Participants received a laptop with calibrated headphones and completed ten 30-minute trainings at home. In the training, participants heard a single sentence each spoken by two concurrent speakers, both the same sentence except for three keywords. The participant is tasked to first identify the correct speaker (using the first keyword) and then recall the remaining keywords via two multiple-choice questions. The less cognitively demanding task must recall the keywords of the current sentence, whereas the more cognitively demanding task must do so for the previous sentence. The SNR of the stimuli changed adaptively based on the users’ performance to achieve an optimal difficulty level for each iteration. To adjust SNR, the attended speaker’s volume was kept constant, and the unattended speaker’s volume varied with the participants’ performance.

### C. Neural Measures

#### 1) Data Collection

The MEG portion of this experiment consisted of participants listening to 60-second excerpts from an audiobook in the public domain (narrated by a male [13] and female [14] speaker) under four conditions: 1) a target speaker with no competing speaker (“quiet”); 2) a target speaker and four competing speakers simulating multi-talker babble (“babble”); 3) a target speaker with the opposite-sex speaker presented at the same loudness (“0 dB SNR”); 4) a target speaker with the opposite-sex speaker presented 6 dB louder than the target speaker (“-6 dB SNR”)

Each noise condition contained separate exemplars for each sex. Each passage was repeated three times, and the listener received verbal and visual instructions as to which sex speaker they should listen to. After each repetition, participants answered a simple yes/no comprehension question designed to keep participants on task; subject-reported intelligibility and listening effort was also logged.

Stimuli were normalized to an equal perceptual loudness and presented diotically using insert earbuds at 70 dB sound pressure level. Subjective listening effort was analyzed by taking the mean across the competing speaker conditions (babble and two-speaker). Neural analysis only included the two-speaker conditions (0 dB and −6 dB SNR) as the power spectrum of the babble condition was different from the two-talker conditions.

#### 2) MEG Preprocessing

Neural data was recorded using a 157-magnetometer whole-head MEG system (Kanazawa Institute of Technology). Data was sampled with a 2 kHz sampling rate and filtered with an online, lowpass filter at 500 Hz. Neural data was further processed offline using Eelbrain (version 0.38) [15] and MNE-Python (version 1.3.0) [16], and denoised with temporal signal-space separation (tSSS) [17]. Data was filtered from 1-40 Hz with a zero-phase FIR filter, and independent component analysis (ICA) was used to remove physiological and noise artifacts. Speech features and neural data were filtered from 1-10 Hz and resampled to 200 Hz.

#### 3) Data Analysis

Neural data was analyzed using encoding and decoding models via the temporal response function (TRF) framework [18], [19]. The encoding and decoding models were used to predict neural responses and reconstruct speech features, respectively, and were derived using the boosting algorithm to obtain a sparse estimation [20].

In the encoding model, model weights represent an impulse response function, called the TRF, that generalizes the time-locked neural response to stimulus features. Suppose we represent the neural response as *y* with time samples *t = 1, …, T* and localized neural sources *n = 1, …, N*. Let *x* represent the matrix of stimulus features where each row *i* contains a different stimulus feature and each column contains the feature values at the time samples *t*. Let *h* represent the model weights where *τ* represents the time lags, occurring from −100 to 700 ms of stimulus onset. Model weights for each neural source are created independently using all stimulus features, where the residual is represented by *ε*, shown in (1).

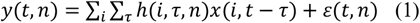

A total of eight speech features (acoustical, lexical, and sub-lexical) were derived from the stimulus including envelope, envelope onset, word onset, unigram word surprisal, contextual word surprisal, phoneme onset, phoneme surprisal, and cohort entropy (see [21] for more details). All eight speech features were used simultaneously to create the encoding models to explain variance, though only the word onset feature encoding model analysis is presented here.

Conversely, in the decoding model, convolution of the neural responses with a weight matrix was used to reconstruct stimulus features, specifically speech envelopes. Denoising source separation (DSS) was used to increase the neural SNR [22]; only the first six DSS components, most coherent across repeated trials, were used—therefore, *m = 1*, ‥, *6*. Let 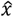 represent the reconstructed speech envelope, and let *g* represent the weight matrix where *τ* represents the time lags, occurring from −500 to 0 ms. The decoding model used all neural responses to contribute to the reconstruction of the speech envelope, shown in (2). The reconstructed speech envelope was correlated with the actual speech envelope to obtain a linear correlation coefficient (r), a measure of reconstruction accuracy.

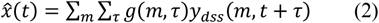

### D. Behavioral Measures

#### 1) Reading Span (RSPAN)

The RSPAN test was used to measure visual working memory [23]. In this task, participants read simple sentences aloud, one word at a time. After each sentence, participants indicated whether the sentence did or did not make sense. After a set of sentences (starting at three and increasing to six), participants were tasked with recalling the final word of each sentence in order. For the present analysis, the total number of correctly recalled final words was used as a measure of working memory.

#### 2) Speech-in-Noise Perception (SPIN)

The bespoke SPIN task was used to measure auditory stream segregation performance. Because the MEG task contains minute-long segments of audio, it is difficult to measure behavioral performance during the MEG task. Instead, a separate SPIN task used short segments of the same story (but unheard) in the various noise conditions; scores are based on the number of keywords (5-7) recalled correctly.

#### 3) Stochastic Figure-Ground (SFG)

The SFG task measures nonlinguistic auditory stream segregation using a group of inharmonic, temporally coherent tones (a “figure”) amongst a background of random tones [24]. Participants must press a button when they believe a figure is present. This task varied randomly in difficulty by changing the number of temporally coherent tones within a “figure” set at 4-, 6-, and 8-chords (called *coherence levels*), with more tones resulting in easier figure detection. Half of the trials contained no figure. Participants were scored based on their accuracy.

### E. Statistical Analysis

Throughout this paper, we use the terms “age” and “session” to distinguish between our comparisons. “Age” refers to comparisons of younger adults and older adults pre-training, and “session” refers to older adults pre- and post-training. Although two similar training groups were implemented, they were combined for statistical analysis due to the small number of participants in each group.

Neural and behavioral data was analyzed using R (version 4.4.1) [25] with linear models (LM), linear mixed effects models (LMEM) [26], or generalized linear mixed effects models (GLMM) [27]. To obtain each model, the buildmer package was used to prune a maximal model and obtain an optimal fit using forward selection and backward elimination [28]. For each model, the performance package was used to verify model assumptions [29]. An ANOVA was performed to compare model fit compared to a simpler model to obtain p-values. If a significant interaction was observed, we performed post hoc testing to obtain the estimated marginal means using the emmeans package [30]. Pairwise comparison p-values were adjusted using the Holm-Bonferroni method.

## III. Results

### A. Age Comparisons

Behavioral analysis showed that younger adults had higher performance scores than older adults across various tasks such as RSPAN (*F* = 47.10, *p* < 0.001) and SPIN (*χ*^2^= 6.82, *p* < 0.01). Interestingly, for the SFG task, a significant interaction of *age* and *coherence level* was found. A post hoc test revealed that this significance was only seen in the easiest 8-chord coherence level where younger adults were more accurate than older adults (est. = 0.505, SE = 0.170, *z* = 3.045, *p* < 0.01) but no differences were found for the harder 4-chord (est. = 0.022, SE = 0.150, *z* = 0.155, *p* = 0.88) or 6-chord (est. = 0.253, SE = 0.150, *z* = 1.685, *p* = 0.18) coherence level.

We analyzed the MEG data from the two-talker noise conditions (0 dB and −6 dB SNR). Initial visual investigation of the stimulus reconstruction accuracies for the two noise conditions revealed large differences in attentional variance between the two age groups. Due to the unequal variances, separate models were created for each age group and no comparisons were performed between the age groups. Fig. 2 shows the predicted reconstruction accuracies of the two models in the two-talker condition. Younger adults have a significant contrast of attention (est. = 0.013, SE = 0.004, *t* = 3.192, *p* < 0.01) that is not seen here in older adults (est. = 0.001, SE = 0.008, *t* = 0.176, *p* = 0.86).

**Figure 1.**
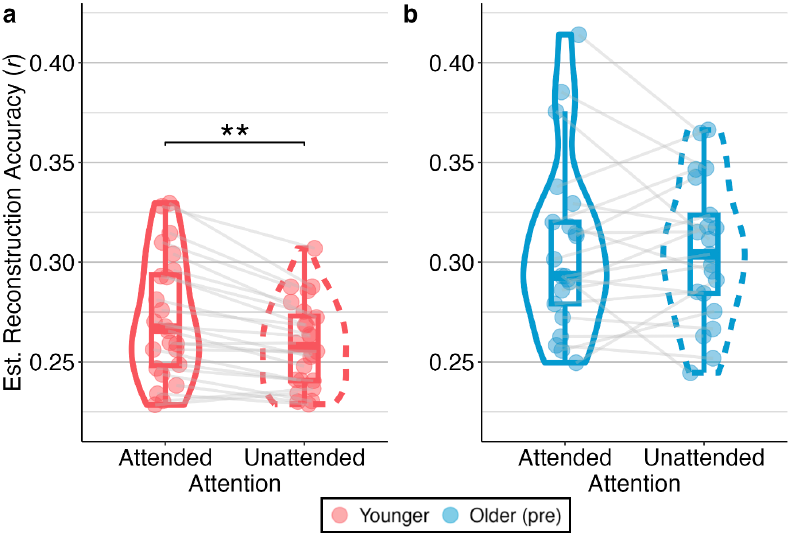
Two separate models predicted values for decoding speech envelopes for attended and unattended speakers in the two-talker noise conditions. (a) Younger adults have higher cortical representations of attended acoustical information than unattended. (b) Older adults may not have these attentional contrasts and may be an indication of speech-in-noise difficulties. (***p < 0.001, **p < 0.01, *p < 0.05)

**Figure 2.**
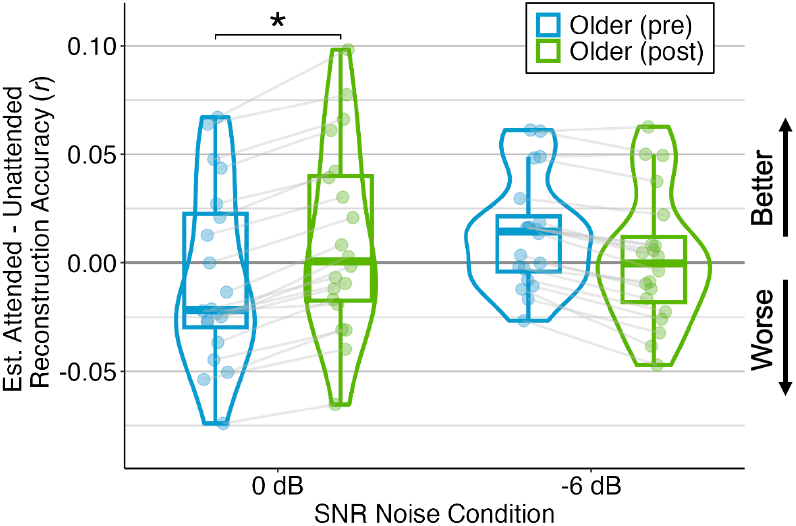
Model-predicted reconstruction accuracies of attended minus unattended speech envelopes in the two-talker MEG noise conditions. Post-training participants showed positive improvement in the 0 dB SNR condition.

### B. Session Comparisons (Pre-vs Post-Training)

Older adults showed improvement in the various behavioral tasks: RSPAN, SPIN, and SFG. The effect of *session* (pre vs. post) was significant for RSPAN (*F* = 15.79, *p* < 0.001), SPIN (*χ*^2^= 8.40, *p* < 0.01), and SFG (*χ*^2^= 6.04, *p* < 0.05). No significant interactions were found between *coherence level* and *session* for the SFG task (*χ*^2^= 4.49, *p* = 0.11).

For subjective ratings, a model of subjective listening effort suggests lower listening effort (*F* = 4.999, *p* < 0.05) after training during the two-talker and babble conditions. While listening effort reports were subjective, neural analysis also supported decreased listening difficulties after training. After training, the peak of the late response (M400) of the word onset TRF, a potential indicator for intelligibility [31], showed differences in the attended speaker (est. = 3.647e-05, SE = 1.456e-05, *t* = 2.504, p < 0.05) and not in the unattended speaker (est. = −7.537e-06, SE = 1.456e-05, *t* = −0.517, *p* = 0.61) in the two-talker noise conditions.

Additionally, a model of speech envelope reconstruction accuracy revealed a significant interaction of *session, attention*, and *SNR*. Fig. 3 shows the outputs of a post hoc analysis, showing greater attentional contrast in the 0 dB SNR condition (est. = −0.018, SE = 0.007, *t* = −2.624, *p* < 0.05); though, the −6 dB condition did not show any attentional difference of *session* (est. = 0.012, SE = 0.007, *t* = 1.766, *p* = 0.08).

**Figure 3.**
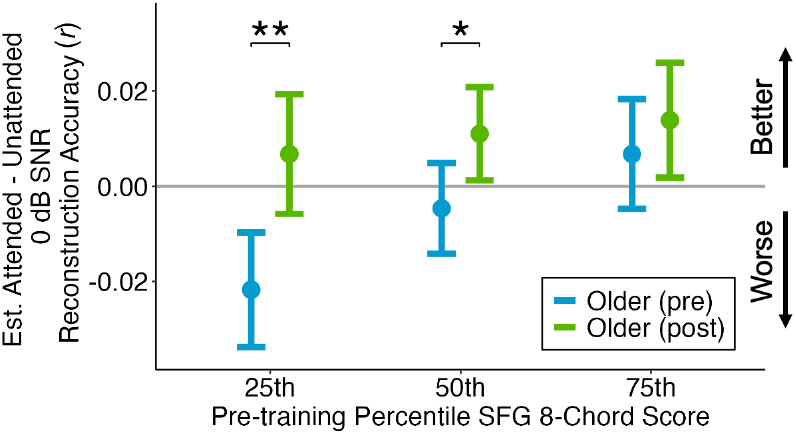
Pre-training performance on SFG (8-chord coherence) as a predictor of change in attentional contrast of reconstruction accuracy in the 0 dB SNR noise condition. Error bars represent the standard error.

A question remaining from this analysis is whether auditory-cognitive training benefits a subset of participants or if this improvement is consistent across all participants. Therefore, we used the reconstruction accuracies of the 0 dB SNR condition with a pre-training behavioral measure to see if neural changes can be predicted. As reported in the previous section, age-related behavioral differences were found in RSPAN, SPIN, and SFG (8-chord). These measures were independently tested to predict neuroplasticity change. Interestingly, there was a significant interaction between performance in the 8-chord coherence level of the SFG task, *SNR*, and *session* (*F* = 7.09, *p* < 0.01). Post hoc testing, shown in Fig. 4, revealed that participants with d-prime in the 25th (est. = −0.025, SE = 0.007, *t* = −3.513, *p* < 0.01) and 50th percentile (est. = −0.018, SE = 0.006, *t* = −2.950, *p* < 0.05) of the SFG 8-chord coherence level had significant attentional differences in their reconstruction accuracies, but participants who scored in the 75th percentile did not show any significant differences after training (est. = −0.018, SE = 0.007, *t* = −1.533, *p* = 0.25).

## IV. Discussion

The main goal of this study was to explore the neural and behavioral impacts of auditory-cognitive training in older adults. We employed encoding and decoding models to observe differences across age groups, guiding our focus on what aspects to explore across training groups. Studies have shown that the reconstructed (decoded) speech envelope of the attended speaker is generally higher than the unattended speaker, as expected [19], [32]. In our study, we observed this effect in younger adults but not in older adults. An equal cortical representation of attended and unattended speech envelopes may suggest that the acoustic information from the unattended speaker is being processed as much as the attended, which may reflect the listening difficulties that older adults have in noisy environments. In contrast, another study has observed an attentional contrast in older adults [12], so the difference between the groups seen here may depend on the specific stimuli and tasks employed.

In the training group comparisons, our behavioral results replicated findings from previous studies that showed improved scores in speech perception and working memory tasks after training [10], [33]. Additionally, subjective listening effort ratings showed a reduction in listening effort for the babble and two-speaker noise conditions. While reports were subjective, neural analysis also supported decreased listening difficulties after training. Specifically, the late peak (M400) of the word onset TRF, a potential indicator for intelligibility [31], showed differences in the attended speaker and not in the unattended speaker post-training. Combining both subjective ratings and objective measures, this provides further support that auditory-cognitive training may improve speech-in-noise performance for older adults who have normal-to-near-normal hearing thresholds.

Neural evidence also showed improved auditory stream segregation in the post-training group. Outputs from the two-speaker decoding models showed larger differences in speech envelope reconstruction accuracies between attended and unattended speakers after training in the 0 dB SNR noise condition. This may be an indication that post-training participants were able to ignore the background speaker more easily.

To see if this change in attentional contrast can be predicted, we then expanded our analysis of the two-talker reconstruction accuracies against a pre-training behavioral measure, the 8-chord coherence SFG task. Our results revealed that older adults who performed worse on this task had the most amount of neuroplasticity change between the attended and unattended speech envelopes. This may suggest that auditory-cognitive training is most effective for participants who demonstrate poor auditory stream segregation, even for a non-speech task, but less effective for participants who are already performing near ceiling.

Overall, the findings of this study provide neural and behavioral evidence for auditory-cognitive training as a method for improving speech-in-noise performance for older adults with normal-to-near-normal hearing thresholds. Neural, subjective, and behavioral measures showed positive improvements. Importantly, older adults who demonstrate worse auditory stream segregation may receive more benefit from auditory-cognitive training.

